# Quantifying Population-level Neural Tuning Functions Using Ricker Wavelets and the Bayesian Bootstrap

**DOI:** 10.1101/2024.05.22.595429

**Authors:** Laura Ahumada, Christian Panitz, Caitlin Traiser, Faith Gilbert, Mingzhou Ding, Andreas Keil

## Abstract

Experience changes the tuning of sensory neurons, including neurons in retinotopic visual cortex, as evident from work in humans and non-human animals. In human observers, visuo-cortical re-tuning has been studied during aversive generalization learning paradigms, in which the similarity of generalization stimuli (GSs) with a conditioned threat cue (CS+) is used to quantify tuning functions. This work utilized pre-defined tuning shapes reflecting prototypical generalization (Gaussian) and sharpening (Difference-of-Gaussians) patterns. This approach may constrain the ways in which re-tuning can be characterized, for example if tuning patterns do not match the prototypical functions or represent a mixture of functions. The present study proposes a flexible and data-driven method for precisely quantifying changes in neural tuning based on the Ricker wavelet function and the Bayesian bootstrap. The method is illustrated using data from a study in which university students (n = 31) performed an aversive generalization learning task. Oriented gray-scale gratings served as CS+ and GSs and a white noise served as the unconditioned stimulus (US). Acquisition and extinction of the aversive contingencies were examined, while steady-state visual event potentials (ssVEP) and alpha-band (8-13 Hz) power were measured from scalp EEG. Results showed that the Ricker wavelet model fitted the ssVEP and alpha-band data well. The pattern of re-tuning in ssVEP amplitude across the stimulus gradient resembled a generalization (Gaussian) shape in acquisition and a sharpening (Difference-of-Gaussian) shape in an extinction phase. As expected, the pattern of re-tuning in alpha-power took the form of a generalization shape in both phases. The Ricker-based approach led to greater Bayes factors and more interpretable results compared to prototypical tuning models. The results highlight the promise of the current method for capturing the precise nature of visuo-cortical tuning functions, unconstrained by the exact implementation of prototypical a-priori models.

**Highlights:**

- Tuning functions are a common way for describing sensory responses, primarily in the visual cortex.
- The quantification and interpretation of tuning functions has faced computational and conceptual problems.
- We demonstrated how the Ricker function can be used as a simple and interpretable way for measuring tuning functions.
- We applied a Ricker function together with a Bayesian Bootstrap approach across a gradient of stimulus features in a generalization conditioning task to characterize visual tuning in the human EEG data.

## Introduction

Neurons in the mammalian primary visual cortex have the capacity to alter their firing pattern as a function of experience, for example in paradigms such as perceptual learning or aversive conditioning (Li et al., 2019). Specifically, neurons that are initially tuned to prefer certain stimulus properties such as a given orientation, location, or spatial frequency may change their tuning function and respond preferentially to a different stimulus property, one that has attained behavioral relevance through learning (Shuler & Bear, 2006). Experience-dependent re-tuning of sensory neurons has been characterized across a range of species, including humans, and from the microscopic level of single-unit recording to the macroscopic level of human brain oscillations (see Li & Keil, 2023 for a review). In animal studies, it has been demonstrated that associative learning sharpens the tuning to preferred stimulus orientations (Gilbert, 1996; Li et al., 2019; Somers et al., 1995; Treue & Trujillo, 1999). This phenomenon has also been referred to as temporal facilitation and occurs at an early stage of visual processing in neurons located in the primary visual cortex (V1; Gilbert, 1996). For instance, the repeated exposure to a visual grating increases the amplitude of the visual evoked potential (VEP) in mouse primary visual cortex (Frenkel et al., 2006).

A large portion of research in the field of sensory re-tuning has relied on aversive Pavlovian conditioning paradigms, in which observers acquire the contingencies between a conditioned threat cue (the CS+) and an unconditioned aversive stimulus (US), such as a loud noise or an electric shock, that is predicted by the CS+ (Ahrens et al., 2015; Li et al., 2019; Moratti & Keil, 2005; Wieser et al., 2014). After acquisition, conditioned aversive responses can become extinguished through a manipulation known as extinction learning, in which the CS+ is repeatedly presented without the US. In differential aversive conditioning paradigms, safety stimuli (CS-), that differ perceptually from the CS+ and are never paired with the US, are employed. For example, the orientation tuning of macaque monkey V1 neurons to rotated Gabor patches (gray-and-white grating stimuli) changed after monkeys had undergone aversive conditioning. More specifically, neurons were tuned to maximize firing in response to the CS+ orientation, and this change in visuo-cortical tuning was specific to the visual field where the CS+ orientation was presented, suggesting a primary visual locus of plastic changes (Li et al., 2019).

In human studies of aversive conditioning, experience-dependent re-tuning can be investigated at the population level by measuring the large-scale neuronal activity in occipito-parietal areas using electroencephalography (EEG). For example, the C1, an early component of the EEG-derived visual event-related potential (ERP) that occurs 40-90 ms post-stimulus increases in amplitude when observers view cues that predict an aversive outcome (Skrandies & Jedynak, 2000; Stolarova et al., 2006; Thigpen et al., 2017). The steady state visual evoked potential (ssVEP) is another EEG-derived population response, which is evoked by stimuli that are periodically modulated in contrast or luminance (Wieser & Keil, 2011). Mirroring findings with the C1 component, a ssVEP evoked by an aversive CS+ is enhanced during conditioning, whereas an ssVEP evoked by a CS-remains unchanged (Miskovic & Keil, 2012; Moratti et al., 2006). A third approach is measuring the event-related change in the amplitude of brain oscillations. Using this approach, it has been demonstrated that aversive conditioning prompts a reduction of power in the alpha frequency range (8-13 Hz) that is stronger for the CS+ than for the CS-(Panitz et al., 2019, Keil & Friedl, 2021, Bacigalupo & Luck 2022).

The re-tuning of visuo-cortical neurons during aversive conditioning can be studied by evaluating the visuocortical responses to a series of non-paired cues that gradually vary in similarity with the CS+, e.g., differing in orientation, size, or color (Lissek et al., 2008). This variation of aversive Pavlovian conditioning is also referred to as aversive generalization learning. It allows researchers to quantify the effect of aversive conditioning on the tuning across a generalization gradient that is formed by the CS+ and the non-paired generalization stimuli (GSs). Previous studies using aversive generalization learning have shown that the resulting re-tuning of visuo-cortical population responses to the critical feature may be characterized by two broadly defined, prototypical, types of tuning patterns (see Figure 1), generalization and sharpening (e.g., McTeague et al. 2015).

Sharpened re-tuning is characterized by heightened responses to the CS+, accompanied by a maximal suppression of responses to the GSs that are most similar to the CS+ (Antov et al., 2020; McTeague et al., 2015; Stegmann et al., 2020). This type of re-tuning has been hypothesized to result from suppressive interactions between neighboring neuronal populations tuned to similar feature levels, such as the sharpening between adjacent columns of orientation/direction specific V1 neurons through lateral inhibition (Crook et al., 1998; McTeague et al., 2015). Generalization re-tuning is also characterized by heightened responses to the CS+, but accompanied by a gradual decrease in GS-evoked responses as a function of decreasing similarity with the CS+. This tuning pattern has been observed repeatedly in fMRI-BOLD responses from brain regions that are sensitive to aversive stimulation. This includes the anterior insula (Lissek et al. 2014), but also fronto-parietal cortical regions associated with attention control (McTeague et al., 2015). It has also been found in selective reduction of alpha-band power evoked by CS+ and GS (Friedl & Keil, 2020, 2021). This phenomenon is often attributed to generalization mechanisms in perception, attention, and motor preparation, in which cue-evoked responses are proportional to the similarity of the cue with a task-relevant stimulus (Martinez-Trujillo & Treue, 2004).

Beyond these broad categories of re-tuning patterns, visuo-cortical tuning has been found to vary with a variety of internal and/or external factors such as individual differences (Stegmann et al., 2020), the type of the task (Farkas et al., 2023), and the feature along which the CS-similarity is being varied (Friedl and Keil, 2021). Prior research has quantified generalization and sharpening either by means of many pairwise comparisons across the GS-evoked brain responses, or by fitting sets of a-priori weights matching the respective gradient patterns. However, using fixed a-priori models of generalization and sharpening often fails to capture the precise nature of the malleability of population responses under consideration given the variability in re-tuning patterns. For example, a tuning function may have a steep drop-off from the CS+ without showing lateral suppressive effects, showing sub-optimal fits with either a-priori function. Thus, it is desirable to establish a flexible and data-driven method capable of robustly capturing a wider range of possible re-tuning patterns. The present study addressed this problem by proposing a simple approach that uses the flexible properties of the difference of Gaussians functions, or the Ricker wavelet function. This function has only one free parameter, the choice of which allows fitting—and thus characterizing—a wide range of tuning patterns. We propose to combine this function fitting method with a statistical approach that relies on the Bayesian bootstrap, used to quantify the goodness of fit of the Ricker model with a given data set.

We tested the proposed approach on data from a visual-aversive generalization learning experiment with black-and-white circular gratings differing in orientation (1 CS+ and 3 GSs) flickering at 15 Hz to evoke ssVEPs. A loud, white noise burst served as the auditory US. The experimental design consisted of three phases: habituation, acquisition, and extinction. Only the acquisition and extinction phases were analyzed in this study because visuo-cortical re-tuning is expected to arise during these phases. The ssVEP and alpha-band power served as variables of interest. Ratings of US expectancy, valence, and arousal were also collected and used to establish the extent to which participants learned the contingencies during the experiment. The present study contrasted the performance of the Ricker model against established a-priori models (generalization and sharpening, see Figure 1). This allows us to compare the model support for each pattern, comparing statistical outcomes with the standard approach and the proposed approach.

**Figure 1.**
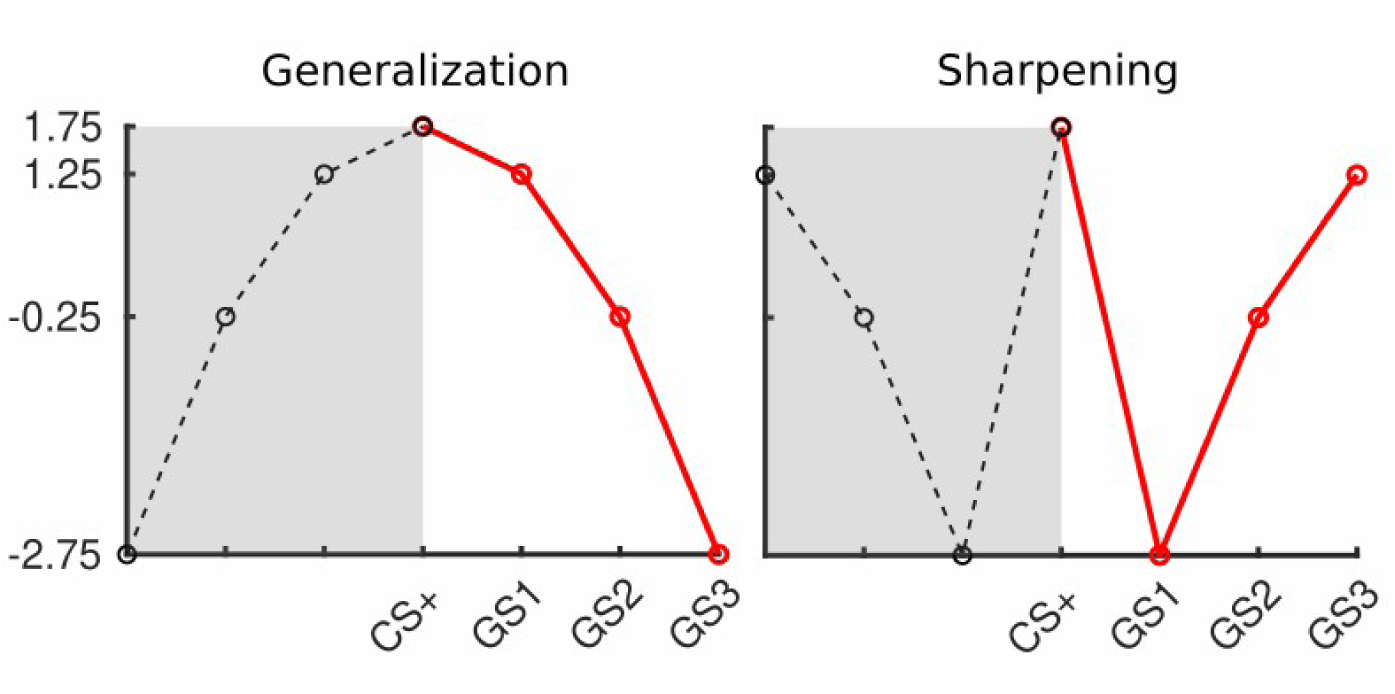
A priori models: generalization weights (1.75,1.25, -0.25, -2.75) and sharpening weights (1.75, -2.75, -0.25, 1.25). The four conditions in the current study covered the right half of the complete model (red lines).

## Material and Methods

### Participants

A total of 37 students (26 female and 11 male) from the University of Florida, (20 white, 2 black/African American, 10 Hispanic, 5 Asian), ranging in age from 18 to 26 years of age (*M* = 19.19) took part in the study. Six participants (4 female and 2 male; 2 white, 3 Hispanic, and 1 Asian) were excluded because of poor EEG signal quality, defined as having Hilbert transformed ssVEP envelopes with a signal-to-noise ratio that was smaller than 2.

Once the participants arrived at the laboratory, they were given an overview of the study, followed by reading and signing the informed consent. All participants reported having no family history of epilepsy and all confirmed having normal or corrected-to-normal vision. After the experimental session, they were compensated with course credits. The study was conducted in accordance with the Declaration of Helsinki and was approved by the University of Florida’s institutional review board.

### Stimuli

White noise with a sound pressure level of 88 or 91 dB(A), see below for details, and a duration of 1000 ms was used as the unconditioned aversive stimulus (US). The noise was delivered through two studio monitor speakers (Behringer Studio 50) located directly behind the participant. High-contrast (100% Michelson contrast) black-and-white circular gratings served as the conditioned stimuli and were occasionally paired (CS+) or never paired (Generalization Stimuli, GSs) with the US. These stimuli varied in terms of the grating orientation (15°, 35°, 55°, and 75°, relative to vertical, see Figure 2a), spanning a similarity gradient for the generalization conditioning paradigm. The four conditions of the study (CS+, GS1, GS2. GS3) were defined by first assigning 15° or 75° as the CS+ or as GS3 (counterbalanced across participants), and the remaining gratings as GS1 and GS2, based on similarity with the CS+. The gratings were presented for 3000 ms at the center of a medium-gray (RGB:127, 127, 127; 45 cd/m^2^) background on a Display++ LCD Monitor (Cambridge Research Systems Ltd., Rochester, UK) with a screen refresh rate of 120 Hz. During US-paired CS+ trials, the final second of the CS+ was accompanied by the US. In order to evoke ssVEPs, gratings were turned on-and-off against the gray background at a constant temporal rate of 15 Hz (i.e., luminance modulation). Participants were seated in front of the screen at a distance of 125 cm measured from the nasion to the middle of the screen, resulting in gratings spanning a vertical and horizontal visual angle of 5°. The background luminance in the chamber was 0.06 cd/m2, as was the black portion of the gratings. The experimental procedure, as well as all auditory and visual stimuli, were coded in Matlab, using Psychtoolbox (Brainard, 1997; Kleiner et al., 2007; Pelli, 1997).

### Aversive generalization learning paradigm

The aversive generalization learning paradigm consisted of three phases (habituation, acquisition, and extinction, see Figure 2b). In the habituation phase, participants saw each of the gratings 8 times (32 trials in total). Ratings of valence, arousal, and expectancy were taken at the end of this initial phase. Valence and arousal were assessed using a scale from 1 to 9, where 1 indicated the stimulus were highly unpleasant or low arousing and 9 indicated it was highly pleasant or high arousing. Expectancy was measured as the probability of occurrence of the US given an specific grating. During the subsequent acquisition phase, the CS+ gratings (15° or 75°) co-terminated with the US for the last 1000 ms of their presentation. The first 6 CS+ trials were always ended with US presentation (i.e. booster trials to facilitate learning), while in the remaining 18 trials the pairing occurred at 50% contingency. The remaining GS gratings were presented 24 times each (total of 96 presentations) and were never paired with the US. This phase was divided into two blocks, so that ratings could be taken in the middle and at the end of the acquisition process. Finally, extinction consisted of the presentation of all four gratings, in 12 trials each, without the US (48 trials in total), and ratings of all stimuli were taken again at the end of this phase. The inter-trial interval in each phase varied between 6500 and 9100 ms (mean = 7000 ms). Breaks were given between phases, which allowed participants to rest as for as long as needed. Then, participants initiated the next phase of the experiment by clicking the mouse. Importantly, participants were not given any explicit information regarding the contingencies between the CS+ and the US; as such, the learning was uninstructed in nature.

**Figure 2.**
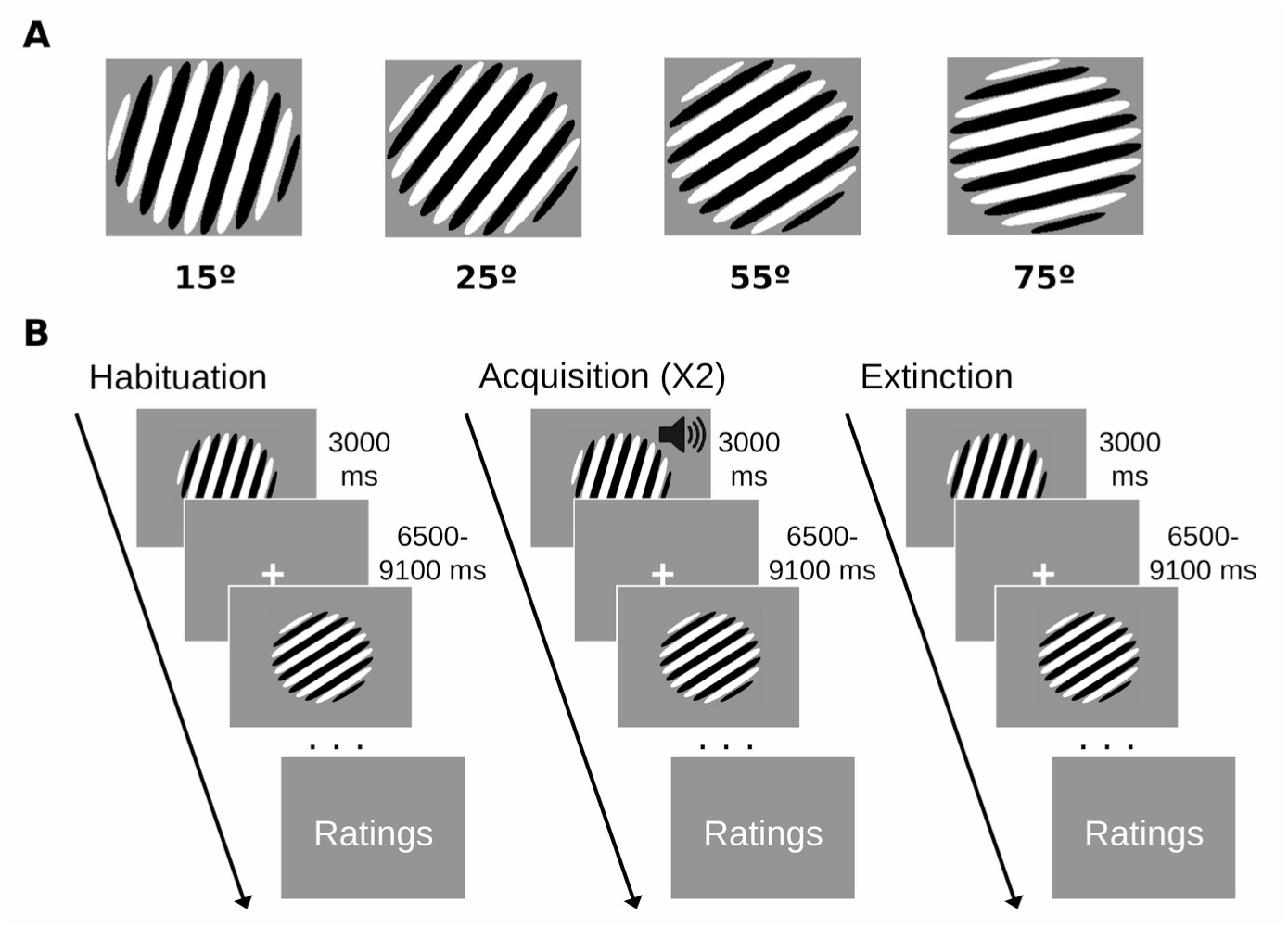
Experimental conditions: grating orientations were assigned to the CS+, GS1, GS2, and GS3 conditions, respectively. 15° and 75° orientation gratings were randomly assigned as the CS+ across participants (A). Experimental design: habituation phase, acquisition phase (divided in two section to get the ratings in the middle and at the end), and extinction phase (B).

### Procedure

Once participants read and signed the consent form, the researcher explained the nature of the study. Then, participants were guided to the experimental chamber. An initial sound test was conducted where the US was presented at 91 dB(A) to determine if participants would tolerate this pressure level. If they indicated that they would not be able to do so, the US was set at 88 dB(A) and used at that level throughout. If participants could not tolerate either of the two sound pressure levels, they were thanked and given class-credits. After this initial test, participants filled out questionnaires targeting symptoms of emotion psychopathology such as anxiety, worry, and depression, not discussed in the present manuscript. Experimenters then fitted the sensor net and prepared the recording. As participants were seated in front of the screen, they were instructed to maintain fixation in the middle of the screen throughout and to avoid body movements during the task. After the experiment, the sensor net was removed and rinsed, and participants were thanked for their participation.

### EEG recording and pre-processing

EEG was recorded using 128-channel HydroCel Geodesic sensor nets and a Net Amp 300 amplifier (Magstim EGI, Oregon, US), with a sample rate of 500 Hz, referenced against Cz. Impedances were kept below 60 KOhms where possible. Subsequently, Emegs2.8 (Peyk et al., 2011) was used for data pre-processing in MATLAB (version 2022, Natick, MA, USA) and the following steps were implemented (Junghofer et al., 2000): the signal was first filtered offline (Highpass = 5 Hz, Lowpass = 40 Hz, using Butterworth filters - 3^rd^ order and 23^rd^ order respectively - with cut-offs defined as 3-dB points). Next, eye blink and eye movement corrections were applied using the regression-based method by Schlögl and colleagues (Schlögl et al., 2007). Data were then segmented from 1000 ms before to 2000 ms after grating onset, resulting in channel by time by trial arrays. Artifact-affected channels and trials were then identified by considering their distribution (across trials and channels) on three quality indices: the median absolute voltage amplitude, the standard deviation of the voltage, and the largest transient value in a given trial and channel. Then, channels that were in the tails of the resulting quality index distributions throughout the recording session were interpolated using spherical splines (maximum of 18 channels, mean = 9.3). Later, trials with artifacts were identified and bad channels interpolated where appropriate: trials in which too many channels would have to be interpolated (defined as the deviation between a forward modeled reference topography with the full channel set and the interpolated channels set,(Junghofer et al., 2000). The average final trials per phase and per condition were: habituation (CS+ = 6.87, GS1 = 7.42, GS2 = 7.03, GS3 = 7.03), acquisition (CS+ = 17.68, GS1 = 18.39, GS2 = 18.42, GS3 = 18.65), and extinction (CS+ = 8.61, GS1 = 8.48, GS2 = 9.39, GS3 = 8.84). Finally, the artifact-free data were re-referenced to the average.

### ssVEP analysis

First, a current source density (CSD) filter was applied to the artifact free trials, to estimate the current density at the surface on the brain, reduce effects of volume conduction, and facilitate interpretability of the topographical information. To this end, the algorithm proposed by Junghöfer and colleagues (Junghöfer et al., 1997) was used with a regularization factor of 5%. From each trial, spectral estimates of the ssVEP amplitude were obtained using a Discrete Fourier Transform (DFT) for each artifact-free trial, applied to the post-stimulus EEG segment starting at grating onset, until 2000 ms post-onset, avoiding the US time period in paired US-CS+ trials. No windowing was applied, as recommended by Bach and Meigen (Bach & Meigen, 1999). The ssVEP amplitude was defined as the DFT amplitude at 15Hz, averaged across the single trial spectra, measured at each electrode.

### Alpha-band power

A Morlet wavelet convolution was conducted to obtain the time-frequency information of the frequency bands between 3.33 and 39.63 Hz (the frequency resolution was 0.33). Specifically, each artifact free trial was convolved with a family of Morlet wavelets with a Morlet coefficient of 10, optimal for analyzing frequencies in the range below 30 Hz. At 10 Hz, these settings resulted in a frequency uncertainty (smearing) of 1 Hz, and a temporal uncertainty of 159 ms (Keil et al., 2022). Then, the resulting single-trial time-frequency representations were averaged and a baseline correction was applied to the result by dividing the average time-frequency matrix by the average power of the baseline segment between -801 and -321 ms. The outcome was then re-scaled to percentage change relative to baseline and averaged across discrete frequencies between 9.33 Hz and 11.33 Hz. Finally, for all statistical analyses, alpha-band power was averaged across a time window from the time point where alpha power decrease was at maximum to the time point where alpha started to increase in all conditions (i.e., 500-1300 ms). This frequency and time range is consistent with prior work on alpha-band reduction in aversive conditioning (Friedl & Keil, 2020, 2021). Besides, given that alpha-band reduction is expressed as a decrement in power, before the statistical analysis, the empirical data was inverted so positive values indicate alpha power reduction.

### Model fitting and statistical analysis

The goal of the present paper was to examine the usefulness of the Ricker function as a flexible and quantitative way to characterize the shape of a generalization gradient with one free parameter. To this end, we applied the Ricker function to the dependent variables formed for the ssVEP and the alpha-band power. We compared the Ricker-based approach to a standard method using a-priori weights. As in prior work (e.g., Stegmann et al., 2020), these weights were multiplied with the empirical data using the dot-product to assess similarity between model-predicted shapes and the empirical shape of the generalization gradient. All statistical analyses were performed using MATLAB (version 2022, Natick, MA, USA). To evaluate aversive learning, changes in alpha-band power and ssVEP amplitude in the four conditions (CS+, GS1, GS2, and GS3) were assessed at all sensors, but focusing on the occipital-parietal region.

*Fitting the Ricker function*. The Ricker wavelet represents a simple mathematical function that reflects the second derivative of a Gaussian, with two parameters, the mean, and the standard deviation. Alternatively, it can be thought of as a difference of two Gaussians, with the standard deviation reflecting the difference in width of the original Gaussians. Here, as in many applications, the mean of the Ricker function is set to 0, and the standard deviation is the only parameter of interest. Its structure is shown in equation 1, below.

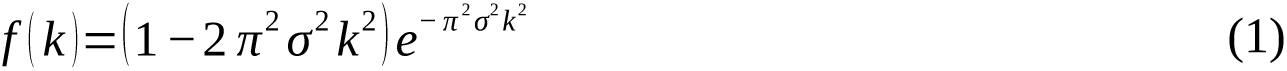

As implemented in the present study, the input into the function is the standard deviation parameter, σ, and the condition number indices *k*, representing the conditions of interest, along the generalization gradient, i.e. CS+, GS1, GS2, and GS3. To heighten the interpretability of the resulting fitted standard deviation (σ) parameters, we chose the vector *k* [0.25 0.50 0.75 1.0] to represent the stimulus gradient with conditions labeled as *k*_1_ (0.25 in the example) to k_n_ (1.0 in the example).

Given equation (1) above, the Ricker function’s shape across its entire range is always symmetrical and is determined by the interaction of the values in *k* as well as the parameter σ. For interpretability of the best-fitting Ricker parameter in the context of tuning functions, one key criterion is the transition from a Gaussian pattern, seen with smaller values of σ, to a Difference-of-Gaussian pattern, generated by greater σ, given *k*. This transition can be readily approximated by the point at which the Ricker function assumes similar values at the two last points in *k* (*k*_n-1_ and *k*_n_). The corresponding critical σ_crit_ is then approximated by the simple equation (equation 2), which finds the σ at which the minimum of the function lies between *k*_n_ and *k*_n-1_ (see Figure 3).

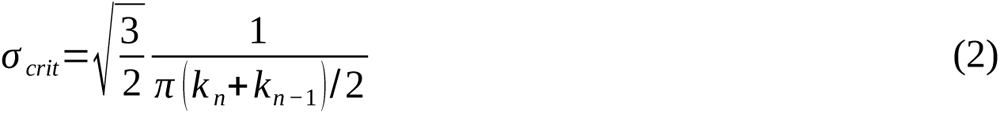

The above equation holds for all monotonically increasing, evenly spaced, vectors *k*. In the present study, our critical σ is estimated as sqrt(3/2) * 1/(pi *(0.75 + 1.0)/2) = 0.45. For researchers studying tuning functions across 9 levels with a *k* vector ranging from 0 to 0.8 in steps of 0.1, for instance, the critical σ would be sqrt(3/2) * 1/(pi * (0.7+0.8)/2) = 0.52.

**Figure 3.**
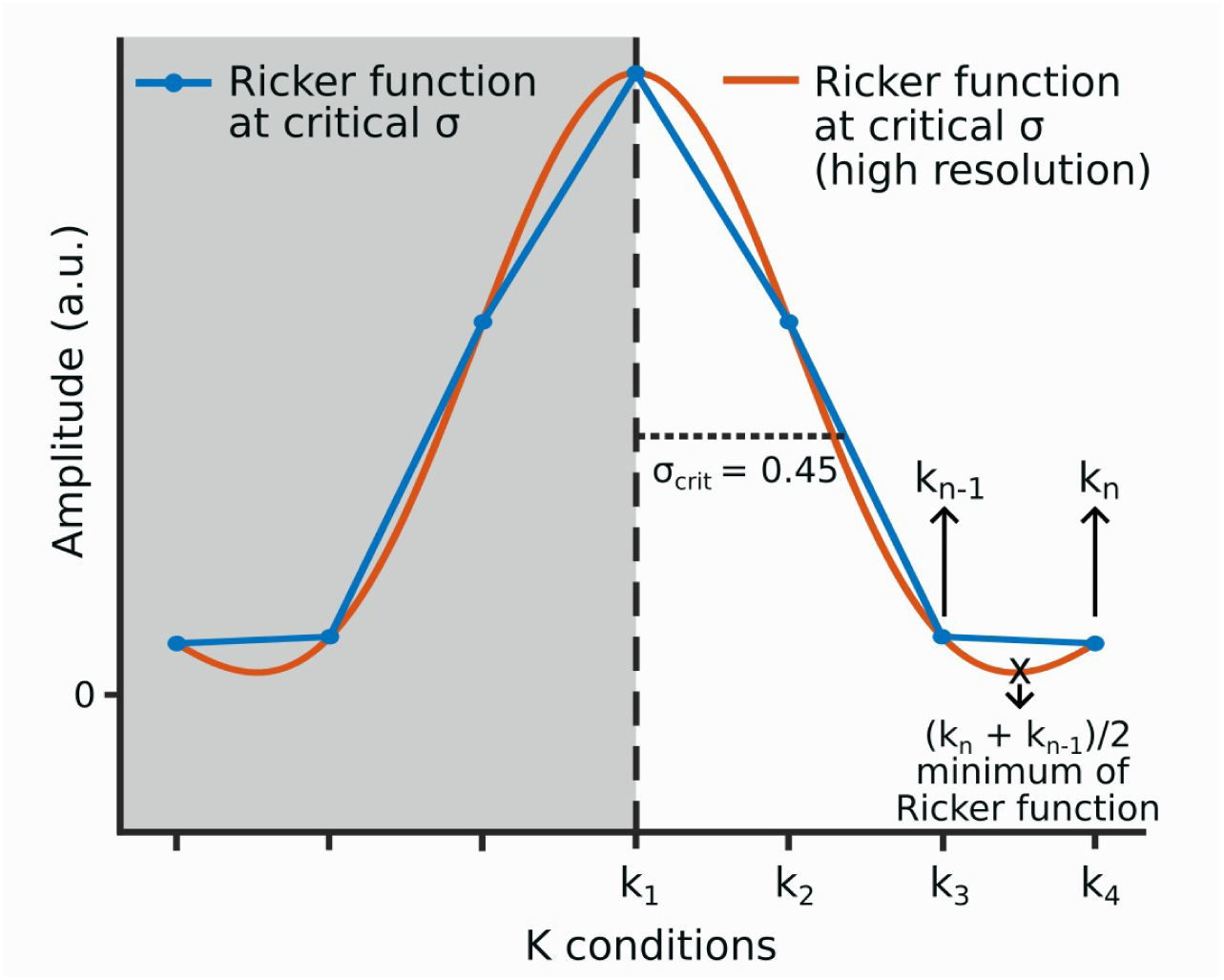
Visual representation of the transition point from a Gaussian pattern to a Difference-of-Gaussian pattern, given a critical value of σ (σ_crit_ = 0.45, for this experiment *k* conditions). The critical σ is determined by finding the minimum of the Ricker function between the two last *k* when they have similar values (*k_n_* + *k_n-1_*)/2).

For each electrode location in the ssVEP and alpha-band data, the Ricker function was fitted using the nonlinear regression function implemented in MATLAB software (nlinfit) with its one free parameter (σ) set to vary freely. This function uses iterative gradient descent with the Levenberg-Marquardt algorithm to find the best-fitting parameter σ, given the data. The termination tolerance for the function value was set to 10^-8^. A maximum of 100,000 iterations was run. The starting value for the standard deviation parameter σ was set to 0.1. The best-fitting parameter σ was used for statistical analyses.

*Fitting the a-priori weights.* To evaluate the fit of the data with the a-priori weights used in prior research (generalization and sharpening weights), dot products between the data and the model weights were computed. Applied to the weights and data from this study, the dot product can be thought of as measure of similarity of correspondence, akin to the covariance. The weights for the generalization model were: 1.75, 1.25, -0.25, and -2.75, and the weights for the sharpening model were: 1.75, -2.75, -0.25, and 1.25 (see Figure 1). The generalization and sharpening weights were orthogonal to each other, thus dot products for those models are zero (uncorrelated). The absolute values for each model summed to the same number, 6, so they did not under-or overweight and thus bias the dot product towards one of the models. For the ratings and alpha-band power, only the a-priori generalization model was tested based on previous research that has shown that alpha reduction is better described by a generalization trend, reflecting the fact that there is no known mechanism that would prompt lateral inhibitory interactions in widespread extra-visual networks involved in alpha suppression (Li and Keil, 2023). For the ssVEP, both the generalization and sharpening models were tested.

*Statistical analysis*. All analyses were based on bootstrapped Bayes Factors (Rubin, 1981) in which first posterior distributions were estimated by means of bootstrapping of the data under each model and then compared to each other or a null model. To this end, participants were randomly (with replacement) included in a parameter estimate and the randomization and estimation repeated 5000 times.

For the Ricker function analyses, and before bootstrapping, the ssVEP and alpha power change data (mean baseline-corrected and inverted-sign alpha-band power during the time window of interest) were range corrected (zero-centered and divided by the range), so that the empirical and predicted data were on the same scale. Then, for each iteration of the bootstrap, a random set of participants (31, the sample size) was drawn with replacement, and the values for each condition (CS+, GS1, GS2, GS3) were averaged across the randomly drawn set, followed by fitting the Ricker function at each electrode location. This was done separately for the acquisition and extinction phases of the experiment. For each iteration, the mean squared error (MSE) between the predicted (based on the best-fitting parameter) and the empirical data was obtained. As a comparison, a null model was also computed using the same bootstrap procedure, but using the residuals relative to a null model in which there were no condition differences. Bayes factors (BFs) were obtained as the ratio of posterior odds over prior odds, where prior odds were set to 1 (equal support for the models to be compared).

For the a-priori weights analyses, the empirical ssVEP, the alpha-band power, and the ratings were also bootstrapped before computing dot products. Additionally, the dot product for a null a-priori model was calculated. For that purpose, the order of the conditions was shuffled before the bootstrapping and the null a-priori model was generated by shuffling the weights order of one of the theoretical models (the generalization weights). The above was used inside the bootstrap iterations to obtain the dot products of the null model. For each iteration of the bootstrap, a set of 31 random participants was drawn, with replacement, and the values for each condition were averaged. Subsequently, the dot product between the averaged bootstrapped data (including the null data) and the a-priori weights (including the null model) was obtained at each electrode location and for the acquisition and extinction phases. Similarly to the Ricker wavelet analyses, BFs were obtained contrasting the dot product of the a-priori model against the dot product of the null model.

The BFs for the Ricker model represent an estimation of the goodness of fit. For this reason, the BF_01_ is reported in this manuscript, since it is predicted that the likelihood of MSE in the null model is greater than the likelihood of the MSE in the Ricker model. On the other hand, the BFs of the a-priori models represent the odds of the model of interest being more similar to the empirical data, thus the BF_10_. All BFs obtained from the EEG data were transformed to the Logarithm of 10 to standardize the values and facilitate interpretation. In addition, only for the purpose of visualizing topographies, a constant of 1 was added to ssVEP’s BFs before taking the logarithm, to facilitate appropriate scaling.

## Results

### Ratings

All ratings followed a generalization trend, with a pattern of heightened expectancy, arousal, and displeasure for the CS+ and decreasing values from GS1 to GS3. This pattern emerged at the mid-point of acquisition and persisted throughout the subsequent phases, but as expected was not present at pre-acquisition (Figure 4). Specifically, for US expectancy, and for the arousal and valence associated with the gratings, the pre-acquisition phase did not show evidence in favor of the generalization trend (expectancy [BF_10_ = 0.23], arousal [BF_10_ = 0.34], and valence [BF_10_ = 1.04]). However, all ratings showed strong to decisive evidence in favor of a generalization trend in the middle of acquisition (expectancy [BF_10_ = 465,124,780.83], arousal [BF_10_ = 127.05], and valence [BF_10_ = 84,381.43]), at the end of acquisition (expectancy [BF_10_ = 128,883,136.55], arousal [BF_10_ = 1,066.11], and valence [BF_10_ = 12,080,968.58]), and after the extinction phase (expectancy [BF10 = 8,130.75], arousal [BF10 = 53.92], and valence [BF_10_ = 351,677.02]).

**Figure 4.**
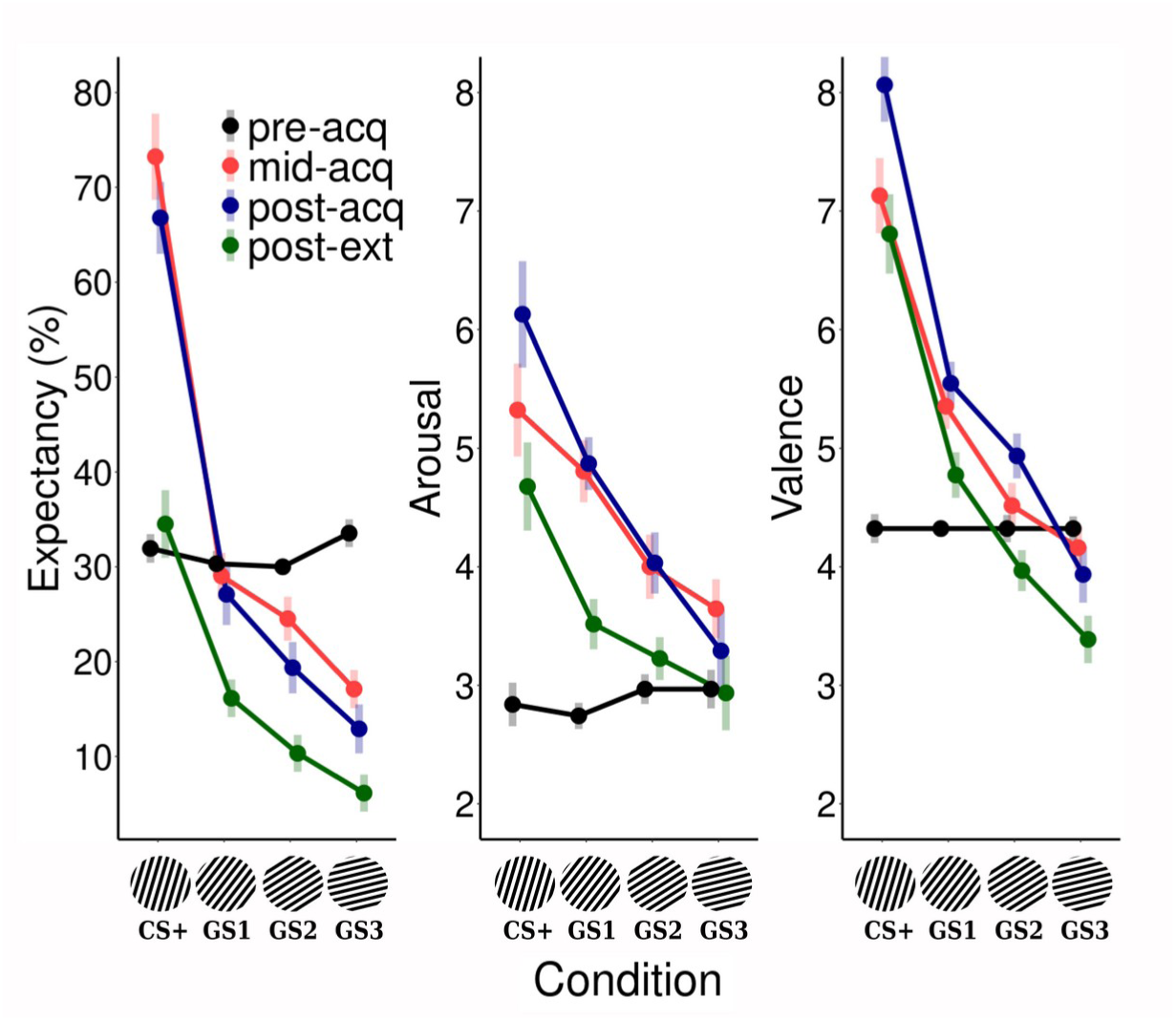
Participants ratings: US expectancy in percentage, arousal, and valence were taken in four different moments throughout the experiment (at the end of habituation, in the middle of acquisition, at the end of acquisition, and at the end of extinction). Error bars correspond to the within-subjects SEM.

### ssVEP

Figure 5 shows the ssVEP in response to the GS1 averaged across all phases, in the time domain (top) and frequency domain (bottom). The topographical plot shows the location of the ssVEP amplitude maximum at the occipital pole.

**Figure 5.**
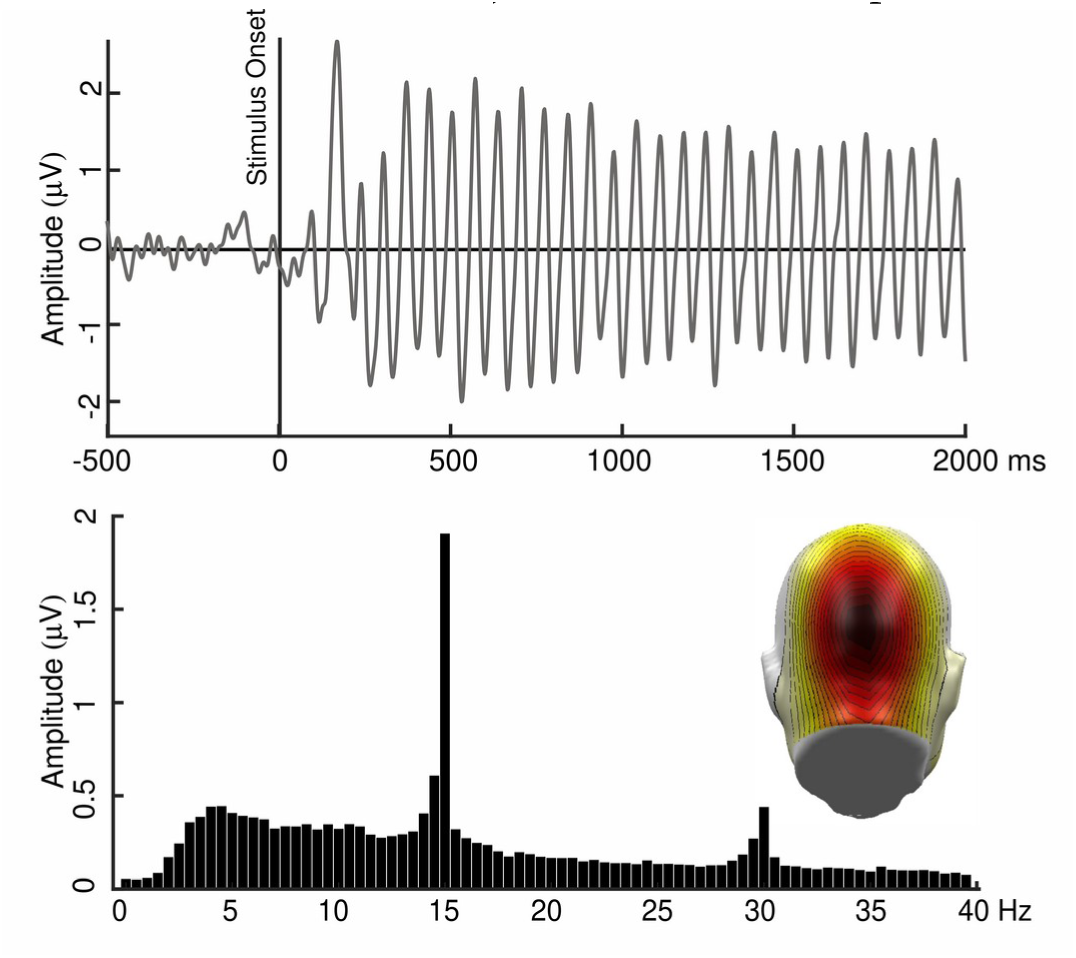
Grand Average ssVEP across phases (GS1 condition). Top panel: ssVEP in the time domain relative to stimulus onset. Bottom panel: Frequency spectrum of the Grand Average with peaks at the 15 Hz and the 30 Hz (2^nd^ harmonic) frequencies, and the topography with the ssVEP peak at the occipital pole.

Figure 6A shows the support for the best-fitting Ricker model relative to the data collected in acquisition and extinction, separately for each sensor. The empirical data and the predicted data are plotted together for each condition and for two sensors that are close to the occipital pole, sensor 74 in acquisition and sensor 75 in extinction. Both sensors are highlighted in Figure 6A. In the acquisition phase, there was decisive support for the hypothesis that the MSE of the Ricker model was smaller than the MSE of a null model at the sensors closest to the occipital pole (74 [LogBF_01_ = 3.097], and 75 [LogBF_01_ = 2.372]). The same results were found in the extinction phase, at the same sensors (74 [LogBF_01_ = 2.397], and 75 [LogBF_01_ = 4.916]).

Regarding the a-priori weights, Figure 6B shows the support for each model relative to the empirical data in acquisition and extinction, again separately computed for each sensor. In acquisition, the generalization weights better described the empirical pattern (given the similarity to the empirical data -dot product-) at the sensors closest to the occipital pole (74 [LogBF_10_ = 1.362], and 75 [LogBF_10_ = 1.001]) than the sharpening weights (74 [LogBF_10_ = 0.091], and 75 [LogBF_10_ = 0.217]). By contrast, in extinction the opposite effect was observed: The sharpening weights showed better fit with the empirical data (74 [LogBF_10_ = 1.834], and 75 [LogBF_10_ = 5.084]), compared to the generalization weights (74 [LogBF_10_ = -0.430], and 75 [LogBF_10_ = -0.360]).

**Figure 6.**
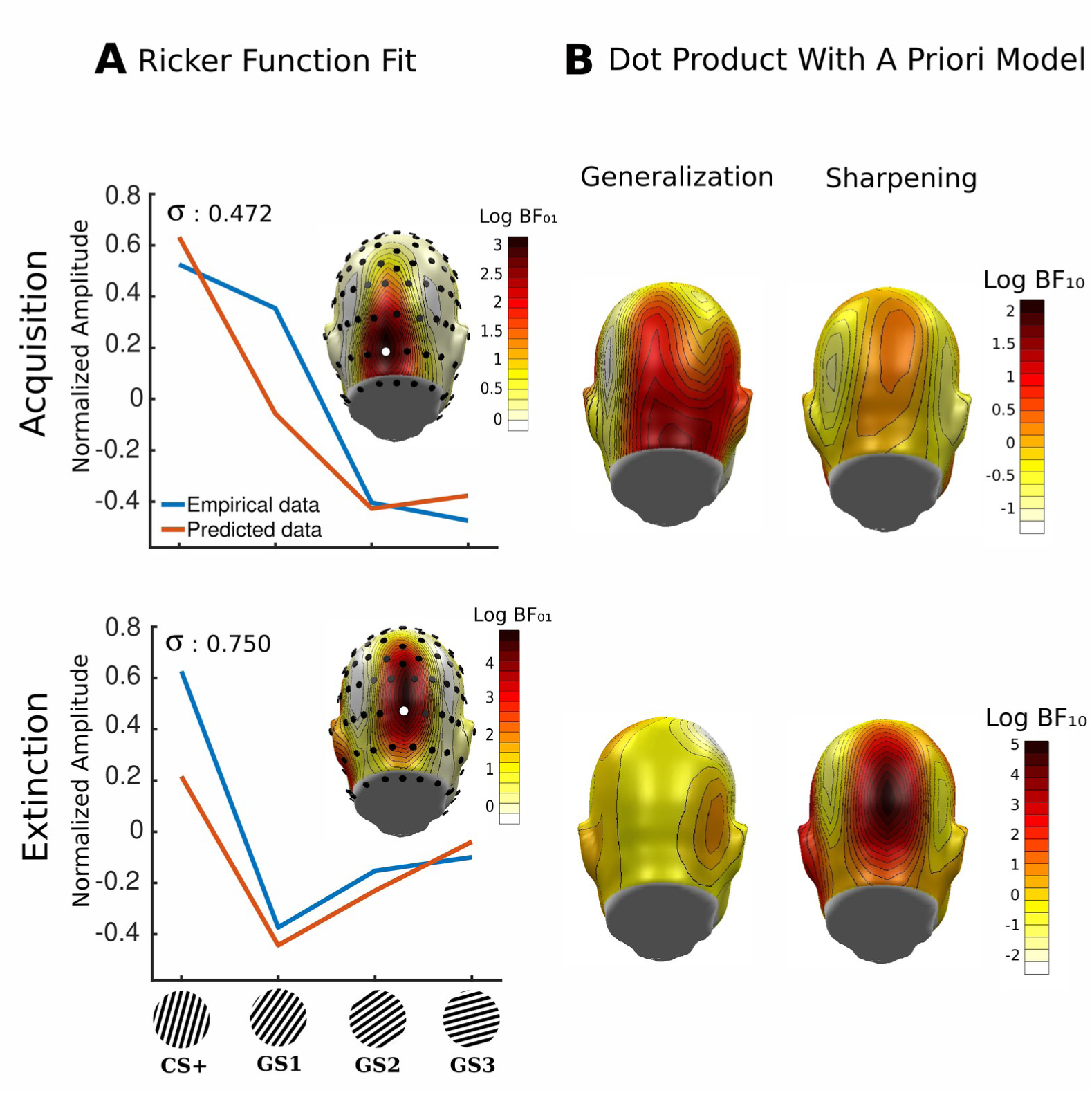
Topographical representation of the logarithmic Bayes Factors from the Ricker model’s MSE against a null model’s MSE in acquisition and extinction for the ssVEP; and the original and predicted data at acquisition and extinction for sensors 74 and 75 (white dots), respectively (A). Topographical representation of the logarithmic Bayes Factors from the comparison of the a priori generalization and sharpening models against a null model in acquisition and extinction (B).

### Alpha-band reduction

The wavelet spectrogram in Figure 7A is an average of the four conditions and sensors in the right parietal region (sensors 77, 78, 84, 85, 90, and 91) and it highlights the reduction of alpha-band power (8 to 13 Hz) in response to the onset of the CS/GS (during the acquisition phase). Figure 7B and 7C show the time course of alpha power for each condition during the acquisition and extinction phases, respectively.

**Figure 7.**
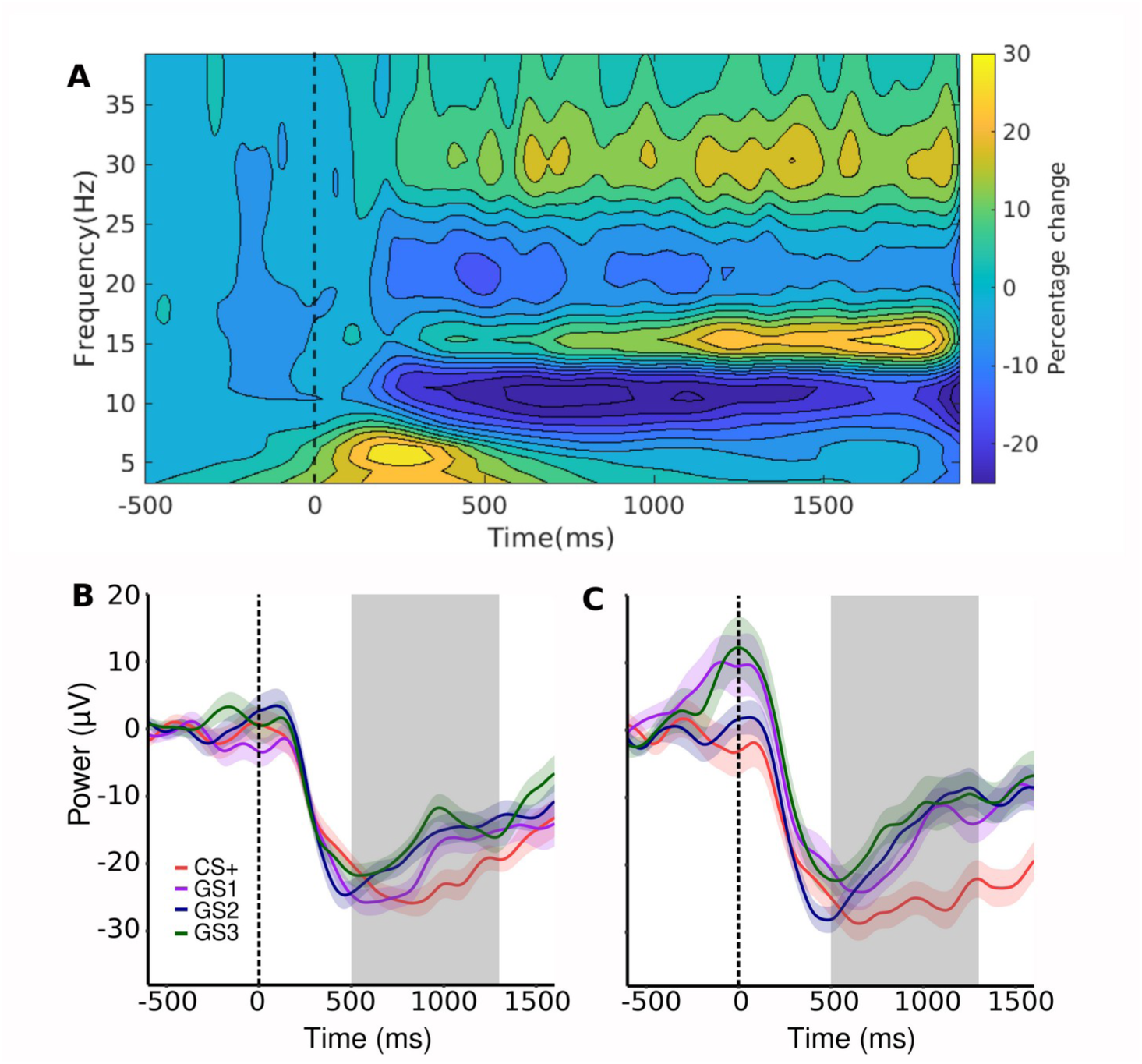
Baseline corrected EEG spectrogram **(A)**. Alpha time course relative to the stimulus onset for the four stimuli during acquisition **(B)** and extinction **(C)**. The gray area represents the time window used for the statistical analyses (500-1300 ms). The line shadows represent the repeated-measure SEM.

The goodness of fit for the Ricker model is shown in Figure 8A for acquisition and extinction on a topographical view. The average inverted values (in the right parietal sensors above mentioned) of each condition for the experimental and predicted data were plotted as well. In general, the Ricker model was well supported by the data at most sensor locations. At sensors around the right parietal region where alpha reduction tended to be most pronounced (77, 78, 84, 85, 90, and 91), the evidence in favor of the Ricker model was decisive in acquisition (mean LogBF_01_ = 2.432) and extinction (mean LogBF_01_ = 2.833). The sensors are highlighted in Figure 8A.

Figure 8B illustrates support for the generalization pattern in acquisition and extinction. There was a strong support, especially at right parietal sensors, for the hypothesis that the generalization trend represented the data in acquisition (77 [LogBF_10_ = 0.961], 78 [LogBF_10_ = 1.353], 84 [LogBF_10_ = 1.422], 85 [LogBF_10_ = 1.534, 90 [LogBF_10_ = 1.237], and 91 [LogBF_10_ = 1.506]) and extinction (77 [LogBF_10_ = 1.403], 78 [LogBF_10_ = 1.140], 84 [LogBF_10_ = 1.606], 85 [LogBF_10_ = 1.490], 90 [LogBF_10_ = 1.679], and 91 [LogBF_10_ = 1.357]).

When subtracting the critical σ_crit_ (0.4502) from the best fitting Ricker σ, the resulting difference map characterizes tuning functions that are more similar to generalization (smaller σ) or difference of Gaussian (greater σ). As shown in Figure 8C, values equal to or lower than zero are more similar to a generalization trend, while values greater than zero are more dissimilar (or more sharpening).

**Figure 8.**
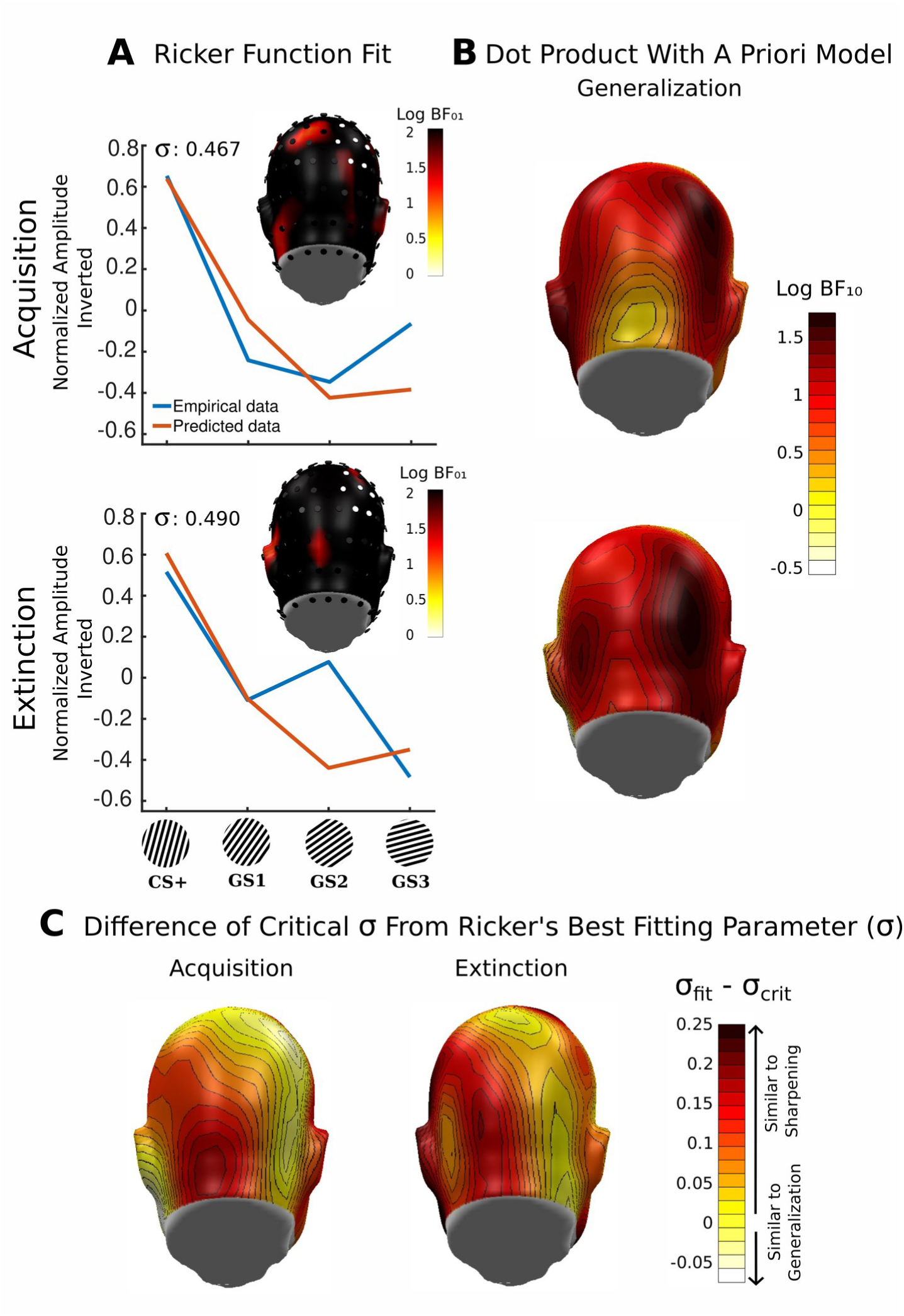
Topographical representation of the logarithmic Bayes Factors from the Ricker model’s MSE against a null model’s MSE in acquisition and extinction for alpha reduction; and the original and predicted data at acquisition and extinction averaged at the right parietal region (sensors -white dots-: 77, 78, 84, 85, 90, and 91) (A). Topographical representation of the logarithmic Bayes Factors from the comparison of the a priori generalization model against a null model in acquisition and extinction (B). Difference between the best fitting parameter and the parameter threshold (σ_crit_, which separates a Ricker model that is more similar to a generalization trend from a Ricker model that is more dissimilar to a generalization trend) for each sensor in acquisition and extinction (C). Values equal or below to zero are more similar to a generalization trend.

## Discussion

The present study aimed to apply a flexible and data-driven mathematical model to evaluate visual-cortical re-tuning during aversive generalization learning, namely the Ricker wavelet model. This model was tested on the ssVEP and alpha-band power. A-priori models of generalization and sharpening were also fit to the data, to compare their goodness-of-fit and sensitivity (expressed as Bayes factors) with fits of the Ricker model. As expected, successful aversive conditioning was reflected in the ratings of the gratings (CS+, GS1, GS2, and GS3) over the experimental session. More specifically, ratings of emotional arousal, hedonic valence, and US expectancy for the CS+ and GSs did not differ before the acquisition phase. However, after the first half of the acquisition phase, the ratings showed a pronounced generalization trend, with heightened ratings for the CS+. This pattern of results was preserved late in acquisition and—to a lesser extent—after extinction, which suggests that extinction was not fully achieved by the end of the experimental session.

### Ricker model fitting of ssVEP amplitude

The application of the Ricker model to the ssVEP data across electrode locations demonstrated that the Ricker model was able to represent the data at sensors located around the occipital pole during acquisition and extinction. This result suggests that the model successfully identified the expected topographical locations, given prior ssVEP research (Wieser et al., 2016). In the same vein, the simple visual grating stimuli used in this study are known to engage lower-tier visuo-cortical neurons at the occipital pole (Verghese et al., 2012), which is in line with the current results. The tuning pattern for the ssVEP amplitude as defined by the best-fitting Ricker model had a generalization-like shape in acquisition and a sharpening-like shape in extinction. These trends corresponded to the a-priori weights (generalization and sharpening weights) that better described the data in acquisition and extinction, respectively.

At variance with prior findings (Antov et al., 2020; McTeague et al., 2015), in the present study the generalization-like trend best characterized the change in the visuo-cortical tuning during acquisition. By contrast, in the above-mentioned studies, the generalization model fit the ssVEP for a short period at the beginning of the acquisition phase. Later in acquisition however, the data were described by a sharpening pattern (Antov et al., 2020; McTeague et al., 2015) as was found during extinction in the present study. A possible explanation for this discrepancy is that in the present study the participants were not informed about the potential relationship between the noise and the gratings (as in Antov et al., 2020, and McTeague et al. 2015). Thus, the emergence of sharpening may have been delayed compared to instructed conditioning, requiring that all of the trials in the acquisition phase occur first, as has been reported in other uninstructed conditioning paradigms (Moratti, Miller, and Keil, 2006).

Another important question in the context of this report is the extent to which prior findings were driven by the use of inflexible, a-priori, models. For example, the generalization weights used in Antov et al. (2020) and McTeague et al. (2015) resembled an over-generalization pattern, the weights of which potentially favoring the lateral inhibition trend, even though the empirical data could have resembled a narrow generalization pattern. Based on the present findings, the characterization of tuning during aversive conditioning may be more precisely and unambiguously defined by the Ricker wavelet model. Specifically, the Ricker model is capable of dynamically and quantitatively define the exact tuning pattern at any given moment in time.

### Ricker fitting and alpha-band reduction

The Ricker wavelet model was able to reflect the patterns of stimulus-induced alpha-band power reduction across the experimental conditions during acquisition and extinction. The topography of model fits was distributed across the scalp, potentially reflecting the fact that alpha-band oscillations in the human EEG tend to spread more widely, compared to the focal ssVEP. However, after subtracting the best fitting σ from the critical σ value (the transition point between a generalization-like and a sharpening-like tuning shape), the alpha reduction pattern was better characterized: the scalp locations showing a generalization-like trend overlapped the locations identified by the a-priori generalization model, namely the right parietal area. Thus, the present Ricker-based approach can be used as an index to define the shape of alpha reduction across sensors.

A generalization of alpha-band power reduction over the generalization gradient in the posterior region is consistent with previous reports by Friedl & Keil (2020, 2021). Over the past decades, researchers have linked the reduction of alpha-band power to active processing of visual information (Klimesch et al., 1998; Mazaheri & Picton, 2005; Pfurtscheller, 2001), while an increase in alpha power has been linked to inhibition of irrelevant information (Klimesch, 2012; Klimesch et al., 2007). Both forms of alpha power changes (power reduction and power increase) are thought to contribute to attentional selectivity during task engagement (Klimesch, 2012). Specifically, during aversive conditioning, the CS+ specific reduction of occipital alpha-band power has been linked to selective attention to the CS+ (Panitz et al., 2019). A generalization of this effect toward similar stimuli has also been found (Friedl & Keil, 2020). Thus, alpha power generalization may index a tuning process of higher-order cognitive processes above and beyond visuo-cortical tuning (Klimesch, 2012). The present study illustrates how tuning processes in higher-order regions and the primary visual cortex can be captured and characterized by a flexible, data-driven approach.

## Conclusions and outlook

Together, the present report suggests that fitting a Ricker wavelet function, in combination with Bayesian bootstrap is a potentially valuable avenue towards quantifying tuning functions obtained from a variety of dependent variables. Specifically, the nature of the Ricker wavelet enables continuous quantification of tuning function properties that does not require a-priori selection of somewhat arbitrary weights. Using such prototypical patterns with specific generalization and/or sharpening weights may obscure and/or mischaracterize meaningful differences in tuning pattern dynamics. This is true especially when tuning patterns show changes that are not readily described by standard Gaussians, or Difference-of-Gaussians. Such changes may involve narrowing of tuning without evidence of lateral inhibitory effects. The overall shape of the best-fitting Ricker wavelet is determined by a single parameter, which covers a wide solution space ranging from a flat Difference-of-Gaussian, to a narrow Difference-of-Gaussian, and ultimately to a prototypical Difference-of-Gaussian with suppressed tails. In this respect, the Ricker wavelet represents a simpler and more readily interpretable approach to fitting tuning functions, compared to more complex functions that have been used for this purpose, for example the Von Mises function (Swindale, 1998). Thus, the best-fitting parameter precisely and gradually characterizes the nature of the empirical tuning profiles on a continuum of possible tuning functions.

In the future, the Ricker wavelet function may also be used to quantify individual differences among participants. For example, the best-fitting Ricker parameter may express the extent to which a given participant shows broadening or sharpening of tuning functions as learning progresses. If reliable and robust, such a parameter could be potentially used to determine relation between tuning dynamics and metrics of inter-individual differences. Furthermore, the Ricker wavelet function can be readily applied to empirical data other than those derived from human EEG signals, including behavioral, autonomic, and hemodynamic brain responses. In summary, the present report illustrates the use of Ricker wavelet fitting in combination with the Bayesian bootstrap, as potentially interesting approach for researchers interested in characterizing tuning functions. Future work will aim to establish the robustness and reliability as well as other psychometric properties of this approach, in larger samples of participants.

## Acknowledgments

The authors would like to thank Richard Ward and Payton Chiasson for assistance in planning and executing the study. This research was supported by grant R01MH125615 from the National Institute of Mental Health.

## References

Ahrens, L. M., Mühlberger, A., Pauli, P., & Wieser, M. J. (2015). Impaired visuocortical discrimination learning of socially conditioned stimuli in social anxiety. Social Cognitive and Affective Neuroscience, 10(7), 929–937. 10.1093/scan/nsu140

Antov, M. I., Plog, E., Bierwirth, P., Keil, A., & Stockhorst, U. (2020). Visuocortical tuning to a threat-related feature persists after extinction and consolidation of conditioned fear. Scientific Reports, 10(1), 3926. 10.1038/s41598-020-60597-z

Bach, M., & Meigen, T. (1999). Do’s and don’ts in Fourier analysis of steady-state potentials. Documenta Ophthalmologica, 99(1), Article 1. 10.1023/A:1002648202420

Bacigalupo, F., & Luck, S. J. (2022). Alpha-band EEG suppression as a neural marker of sustained attentional engagement to conditioned threat stimuli. Social Cognitive and Affective Neuroscience, 17(12), 1101–1117. 10.1093/scan/nsac029

Brainard, D. H. (1997). The Psychophysics Toolbox. Spatial Vision, 10(4), 433–436.

Crook, J. M., Kisvárday, Z. F., & Eysel, U. T. (1998). Evidence for a contribution of lateral inhibition to orientation tuning and direction selectivity in cat visual cortex: Reversible inactivation of functionally characterized sites combined with neuroanatomical tracing techniques. European Journal of Neuroscience, 10(6), 2075. 10.1046/j.1460-9568.1998.00218.x

Frenkel, M. Y., Sawtell, N. B., Diogo, A. C. M., Yoon, B., Neve, R. L., & Bear, M. F. (2006). Instructive Effect of Visual Experience in Mouse Visual Cortex. Neuron, 51(3), 339–349. 10.1016/j.neuron.2006.06.026

Friedl, W. M., & Keil, A. (2020). Effects of Experience on Spatial Frequency Tuning in the Visual System: Behavioral, Visuocortical, and Alpha-band Responses. Journal of Cognitive Neuroscience, 32(6), 1153–1169. 10.1162/jocn_a_01524

Friedl, W. M., & Keil, A. (2021). Aversive Conditioning of Spatial Position Sharpens Neural Population-Level Tuning in Visual Cortex and Selectively Alters Alpha-Band Activity. The Journal of Neuroscience, 41(26), 5723–5733. 10.1523/JNEUROSCI.2889-20.2021

Gilbert, C. D. (1996). Plasticity in visual perception and physiology. Current Opinion in Neurobiology, 6(2), 269–274. 10.1016/S0959-4388(96)80083-3

Junghöfer, M., Elbert, T., Leiderer, P., Berg, P., & Rockstroh, B. (1997). Mapping EEG-potentials on the surface of the brain: A strategy for uncovering cortical sources. Brain Topogr, 9(3), Article 3.

Junghofer, M., Elbert, T., Tucker, D. M., & Rockstroh, B. (2000). Statistical control of artifacts in dense array EEG/MEG studies. Psychophysiology, 37(4), Article 4.

Keil, A., Bernat, E. M., Cohen, M. X., Ding, M., Fabiani, M., Gratton, G., Kappenman, E. S., Maris, E., Mathewson, K. E., Ward, R. T., & Weisz, N. (2022). Recommendations and publication guidelines for studies using frequency domain and time-frequency domain analyses of neural time series. Psychophysiology, 59(5), Article 5. 10.1111/psyp.14052

Kleiner, M., Brainard, D., & Pelli, D. (2007). What’s new in Psychtoolbox*-*3*?*

Klimesch, W. (2012). Alpha-band oscillations, attention, and controlled access to stored information. Trends in Cognitive Sciences, 16(12), 606–617. 10.1016/j.tics.2012.10.007

Klimesch, W., Doppelmayr, M., Russegger, H., Pachinger, T., & Schwaiger, J. (1998). Induced alpha band power changes in the human EEG and attention. Neuroscience Letters, 244(2), 73–76. 10.1016/S0304-3940(98)00122-0

Klimesch, W., Sauseng, P., & Hanslmayr, S. (2007). EEG alpha oscillations: The inhibition–timing hypothesis. Brain Research Reviews, 53(1), 63–88. 10.1016/j.brainresrev.2006.06.003

Li, W., & Keil, A. (2023). Sensing fear: Fast and precise threat evaluation in human sensory cortex. Trends in Cognitive Sciences, 27(4), 341–352. 10.1016/j.tics.2023.01.001

Li, Z., Yan, A., Guo, K., & Li, W. (2019). Fear-Related Signals in the Primary Visual Cortex. Current Biology, 29(23), 4078–4083.e2. 10.1016/j.cub.2019.09.063

Lissek, S., Biggs, A. L., Rabin, S. J., Cornwell, B. R., Alvarez, R. P., Pine, D. S., & Grillon, C. (2008). Generalization of conditioned fear-potentiated startle in humans: Experimental validation and clinical relevance. Behaviour Research and Therapy, 46(5), 678–687. 10.1016/j.brat.2008.02.005

Lissek, S., Bradford, D. E., Alvarez, R. P., Burton, P., Espensen-Sturges, T., Reynolds, R. C., & Grillon, C. (2014). Neural substrates of classically conditioned fear-generalization in humans: A parametric fMRI study. Social Cognitive and Affective Neuroscience, 9(8), 1134–1142. 10.1093/scan/nst096

Martinez-Trujillo, J. C., & Treue, S. (2004). Feature-Based Attention Increases the Selectivity of Population Responses in Primate Visual Cortex. Current Biology, 14(9), 744–751. 10.1016/j.cub.2004.04.028

Mazaheri, A., & Picton, T. W. (2005). EEG spectral dynamics during discrimination of auditory and visual targets. Cognitive Brain Research, 24(1), 81–96. 10.1016/j.cogbrainres.2004.12.013

McTeague, L. M., Gruss, L. F., & Keil, A. (2015). Aversive learning shapes neuronal orientation tuning in human visual cortex. Nature Communications, 6(1), 7823. 10.1038/ncomms8823

Miskovic, V., & Keil, A. (2012). Acquired fears reflected in cortical sensory processing: A review of electrophysiological studies of human classical conditioning. Psychophysiology, 49(9), Article 9. 10.1111/j.1469-8986.2012.01398.x

Moratti, S., & Keil, A. (2005). Cortical activation during Pavlovian fear conditioning depends on heart rate response patterns: An MEG study. Cognitive Brain Research, 25(2), 459–471. 10.1016/j.cogbrainres.2005.07.006

Moratti, S., Keil, A., & Miller, G. A. (2006). Fear but not awareness predicts enhanced sensory processing in fear conditioning. Psychophysiology, 43(2), Article 2.

Panitz, C., Keil, A., & Mueller, E. M. (2019). Extinction-resistant attention to long-term conditioned threat is indexed by selective visuocortical alpha suppression in humans. Scientific Reports, 9(1), Article 1. 10.1038/s41598-019-52315-1

Pelli, D. G. (1997). The VideoToolbox software for visual psychophysics: Transforming numbers into movies. Spatial Vision, 10(4), 437–442.

Peyk, P., De Cesarei, A., & Junghöfer, M. (2011). ElectroMagnetoEncephalography software: Overview and integration with other EEG/MEG toolboxes. Computational Intelligence and Neuroscience, 2011.

Pfurtscheller, G. (2001). Functional brain imaging based on ERD/ERS. Vision Research, 41(10–11), 1257–1260. 10.1016/S0042-6989(00)00235-2

Rubin, D. B. (1981). The bayesian bootstrap. The annals of statistics, 130-134.

Schlögl, A., Keinrath, C., Zimmermann, D., Scherer, R., Leeb, R., & Pfurtscheller, G. (2007). A fully automated correction method of EOG artifacts in EEG recordings. Clinical Neurophysiology, 118(1), 98–104. 10.1016/j.clinph.2006.09.003

Shuler, M. G., & Bear, M. F. (2006). Reward Timing in the Primary Visual Cortex. 311.

Skrandies, W., & Jedynak, A. (2000). Associative learning in humans—Conditioning of sensory-evoked brain activity. Behavioural Brain Research, 107(1–2), 1–8. 10.1016/S0166-4328(99)00096-0

Somers, D., Nelson, S., & Sur, M. (1995). An emergent model of orientation selectivity in cat visual cortical simple cells. The Journal of Neuroscience, 15(8), 5448–5465. 10.1523/JNEUROSCI.15-08-05448.1995

Stegmann, Y., Ahrens, L., Pauli, P., Keil, A., & Wieser, M. J. (2020). Social aversive generalization learning sharpens the tuning of visuocortical neurons to facial identity cues. eLife, 9, e55204. 10.7554/eLife.55204

Stolarova, M., Keil, A., & Moratti, S. (2006). Modulation of the C1 Visual Event-related Component by Conditioned Stimuli: Evidence for Sensory Plasticity in Early Affective Perception. Cerebral Cortex, 16(6), 876–887. 10.1093/cercor/bhj031

Swindale, N. V. (1998). Orientation tuning curves: Empirical description and estimation of parameters. Biological Cybernetics, 78(1), 45–56. 10.1007/s004220050411

Thigpen, N. N., Bartsch, F., & Keil, A. (2017). The malleability of emotional perception: Short-term plasticity in retinotopic neurons accompanies the formation of perceptual biases to threat. Journal of Experimental Psychology: General, 146(4), 464–471. 10.1037/xge0000283

Treue, S., & Trujillo, J. C. M. (1999). Feature-based attention influences motion processing gain in macaque visual cortex. Nature, 399(6736), 575–579. 10.1038/21176

Verghese, P., Kim, Y.-J., & Wade, A. R. (2012). Attention Selects Informative Neural Populations in Human V1. The Journal of Neuroscience, 32(46), Article 46. 10.1523/JNEUROSCI.1174-12.2012

Wieser, M. J., & Keil, A. (2011). Temporal Trade-Off Effects in Sustained Attention: Dynamics in Visual Cortex Predict the Target Detection Performance during Distraction. The Journal of Neuroscience, 31(21), 7784–7790. 10.1523/JNEUROSCI.5632-10.2011

Wieser, M. J., Miskovic, V., & Keil, A. (2016). Steady-state visual evoked potentials as a research tool in social affective neuroscience. Psychophysiology, 53(12), 1763–1775. 10.1111/psyp.12768

Wieser, M. J., Miskovic, V., Rausch, S., & Keil, A. (2014). Different time course of visuocortical signal changes to fear-conditioned faces with direct or averted gaze: A ssVEP study with single-trial analysis. Neuropsychologia, 62, 101–110. 10.1016/j.neuropsychologia.2014.07.009

